# Enterobacterial common antigen biosynthesis in *Yersinia pestis* is tied to antimicrobial peptide resistance

**DOI:** 10.1101/2023.08.26.554945

**Authors:** Kari L. Aoyagi, Basil Mathew, Mark A. Fisher

## Abstract

Resistance to antimicrobial peptides (AMPs) plays an important role in allowing *Yersinia pestis* to maintain a successful infection in the flea vector *Xenopsylla cheopis*. Mutants that are unable to modify lipid A in their outer membrane with aminoarabinose (Ara4N), showed increased sensitivity to AMPs such as polymyxin B (PB), as well as decreased survival in fleas. A deletion mutant of *wecE*, a gene involved in biosynthesis of enterobacterial common antigen (ECA), also displayed hypersusceptibility to PB in vitro. Additional mutants in the ECA biosynthetic pathway were generated, some designed to cause accumulation of intermediate products that sequester undecaprenyl phosphate (Und-P), a lipid carrier that is also used in numerous other pathways, including for peptidoglycan, O-antigen, and Ara4N biosynthesis. Mutants that accumulate Und-PP-linked intermediates (ECA-lipid II) showed increased susceptibility to PB, reduced Ara4N-modified lipid A, altered cell morphology, and decreased ability to maintain flea infections. These effects are consistent with a model where *Y. pestis* has a sufficiently limited free Und-P pool such that sequestration of Und-P as ECA-lipid II prevents adequate Ara4N biosynthesis, ultimately resulting in AMP hypersusceptibility.

## INTRODUCTION

*Yersinia pestis*, the causative agent of plague, undergoes transmission from fleas to mammalian hosts, although much is still unknown about the interaction of the bacteria with their flea hosts. A previous signature-tagged mutagenesis screen performed in a native host, the rat flea *Xenopsylla cheopis*, identified *Y. pestis* KIM6+ mutants with decreased survival in fleas after four days. Most of these mutants also displayed increased sensitivity to antimicrobial peptides (AMPs) such as polymyxin B (PB), suggesting AMP resistance plays an important role in allowing *Y. pestis* to initiate and maintain a successful flea infection [1].

One of the AMP-hypersensitive mutants identified from this screen contained an insertion in the UDP-4-amino-4-deoxy-L-arabinose-oxoglutarate aminotransferase (*arnB*) gene, which is the first gene in the operon involved in aminoarabinose (Ara4N) biosynthesis. Modification of the lipid A portion of lipopolysaccharide (LPS) with Ara4N is a common mechanism known to provide enhanced resistance to AMPs [2, 3]. The *arnB* insertion mutant was more susceptible to the AMPs cecropin A and polymyxin B and produced lipid A lacking Ara4N modifications [1]. However, an in-frame deletion of *arnB* surprisingly retained modest levels of AMP resistance and Ara4N modification, suggesting the presence of a compensatory mechanism. It was determined that WecE, a closely related aminotransferase involved in biosynthesis of enterobacterial common antigen (ECA), could partially offset the loss of ArnB. A double knockout mutant lacking both the *arnB* and *wecE* genes was highly susceptible to PB and contained no discernable amount of Ara4N-modified lipid A detected by MALDI-TOF. However, a clean deletion of *wecE* alone also displayed a hypersensitivity to PB, which suggested a role for ECA in AMP resistance [1].

ECA is a conserved polysaccharide found in nearly all members of the *Enterobacterales* that was described decades ago but its function remains unclear [4]. It is composed of repeating trisaccharides and occurs in several different forms: as a linear polysaccharide linked to phosphatidylglycerol in the outer membrane (ECA_PG_), as an immunogenic linear polysaccharide linked to the LPS core (ECA_LPS_), and as a cyclic polysaccharide localized to the periplasm (ECA_CYC_) [5]. Although ECA is ubiquitous among the *Enterobacterales*, its functions remain obscure due to the intertwined nature of the cell envelope. The ECA biosynthetic pathway shares genes and substrates with other cell envelope pathways, including those for peptidoglycan and O-antigen production, making it difficult to assign specific effects to envelope pathway disruptions [5]. Additionally, the isoprenoid carrier, undecaprenyl phosphate (Und-P), is used by these envelope component pathways, forming polyprenyl-diphosphate intermediates (Und-PP-), as well as other pathways including Ara4N, that form polyprenyl-monophosphate intermediates (Und-P-) [6], so changes in the allocation of Und-P can alter cell envelope composition and structure. Disrupting the ECA pathway at certain points leads to accumulation of Und-PP-linked ECA intermediates that sequester part of the available Und-P pool, leading to alterations in cell size and shape, envelope composition, and envelope stress responses, which can affect cell responses to environmental conditions [7-9]. Here, we describe the characterization of several mutants in the ECA biosynthetic pathway in *Y. pestis* KIM6+.

## MATERIALS AND METHODS

### Bacterial strains and growth conditions

Bacterial strains and plasmids used in this study are listed in Table S1. *Yersinia pestis* strains were routinely grown in Heart Infusion (HI) broth (Difco, BD, Franklin Lakes, NJ) or on HI agar at 28°C. Unless otherwise indicated, antibiotics were used at the following concentrations: 50 μg/ml kanamycin (Km), 100 μg/ml ampicillin (Ap), 5 ug/ml tunicamycin (Tun), and 0.05 ug/ml anhydrotetracycline (ATc).

### Generation and complementation of mutants

Unmarked, in-frame deletions were generated in KIM6+ using the λRed recombineering approach as described previously [1, 18]. Mutant strains were complemented by amplifying the target gene from KIM6+ using primers indicated in Table S2. The resulting amplicons were then cloned into pKLA-KM under control of the *wecA* promoter and mutants were transformed with complementation vectors or empty vector controls by electroporation. Recombinants and complementation vectors were sequenced for verification prior to further experimentation.

### AMP susceptibility assays

AMP susceptibility assays were performed as described by Felek et al. [19] except as otherwise noted. Briefly, two-fold serial dilutions of polymyxin B were prepared in HI broth in 96-well round bottom plates. Bacterial strains were grown overnight at 28°C and cells were added to each well in triplicate at a concentration of 5x10^5 cfu/ml. Plates were incubated 24 h at 28°C and the minimum inhibitory concentration (MIC) for each strain was recorded as the lowest PB concentration at which bacterial growth was not observed. *E. coli* ATCC 25922 was used for quality control.

### Lipid A extraction and MALDI-TOF mass spectrometry

Lipopolysaccharide was extracted from overnight bacterial cultures using an LPS extraction kit (iNtRON Biotechnology, Seoul, Korea) as directed, except 20 ml of culture were initially harvested for use. Lipid A was isolated as described previously [1] and suspended in 50 μl of a chloroform/methanol/water mixture (4:4:1, v/v/v). Lipid A preparations were analyzed by matrix-assisted laser desorption ionization time-of-flight (MALDI-TOF) mass spectrometry in the negative ion mode using a Bruker microflex LRF instrument (Bruker Daltonics, Billerica, MA). Lipid A samples were mixed with an equal volume of 5-chloro-2-mercaptobenzothiazole (CMBT; 20 mg/ml in chloroform/methanol/water [4:4:1, v/v/v]) supplemented with 20 mM EDTA-ammonium hydroxide [20] and 0.5 μl aliquots were spotted onto the target plate. Spectra were acquired in the reflectron mode and analyzed using flexAnalysis (Bruker Daltonics). External calibration was performed using diphosphoryl lipid A from *E. coli* F583 (Sigma-Aldrich, St. Louis, MO).

### ECA extraction and blotting

Two ml of overnight culture was centrifuged, and the resulting pellet was resuspended in 50 μl 0.9% saline to which 284 μl of 100% ethanol was added. Tubes were mixed by inversion and incubated for 5 minutes, then centrifuged again to pellet cellular debris. The supernatant was removed and dried in a SpeedVac. The dried pellet was dissolved in 10-20 μl 1X TBS and 5 μl spots were deposited onto a nitrocellulose membrane and allowed to dry. Membranes were blocked by rocking on a platform for 30 min in blotting buffer (5% [w/v] powdered milk in 1X TBS with 0.1% Tween 20) and this blotting buffer was used for all antibody incubations and washes unless otherwise noted. Membranes were incubated with a 1:1000 dilution of O14 *E. coli* antisera (SSI Diagnostica) for one hour, washed 3 times (5 min ea.), then incubated with a 1:1000 dilution of 0.1 mg/ml phosphatase labeled goat anti-rabbit IgG (KPL) for one hour. Membranes were again washed 3 times, then rinsed once in assay buffer (1 mM Tris, 1 mM MgCl_2_, pH 9.6) and spots were detected with 300 μl BCIP-NBT Phosphatase Substrate (KPL), 10 min.

### Flea coinfection assays

Mutant and wild type *Y. pestis* were added to defibrinated rat blood at a concentration of 10^9^ cfu/ml and *X. cheopis* fleas were allowed to feed using an artificial feeding apparatus [18]. Fleas that did not feed effectively were discarded, while fed fleas were collected and flash frozen at the time of feeding and at 1 and 4 days post-feeding. Thawed fleas were surface sterilized prior to plating by treating with 3% hydrogen peroxide, followed by washes with 95% ethanol and water. Fleas were air dried then crushed in a 1.5 ml microfuge tube containing 200 μl 1X PBS using a sterile disposable pestle. An additional 800 μl 1X PBS was used to rinse the pestle and bring the volume to 1 ml. Flea lysate was diluted 10-fold and 10-20 μl of lysate was plated to non-selective BHI and BHI containing Km using an overlay method described previously [18]. Plates were incubated at 28°C for 3 days before colony counting. Four individual fleas were plated for each coinfection timepoint and ratios of mutant to all cells were calculated.

## RESULTS

### Mutant generation and ECA production

We generated in-frame deletion mutants at different points in the ECA biosynthetic pathway using lambda-Red-based recombineering [10] to better understand the relationship between ECA genes and AMP resistance. These mutants included Δ*wecA*, Δ*wecB*, Δ*wecF* and Δ*wecOP* (an operon mutant lacking all genes from *wecB-G*), in addition to the previously generated Δ*wecE* (Fig. 1). WecA initiates the biosynthesis of ECA and is also involved in O-antigen biosynthesis; however, *Y. pestis* does not produce O-antigen so the Δ*wecA* mutant should not affect other pathways beyond ECA production. Loss of the *wecA* or *wecB* gene should not lead to the buildup of any carrier-linked intermediates, while the loss of *wecE* or *wecF* leads to ECA-lipid II accumulation and sequestration of Und-P (Fig.1), allowing distinction between phenotypes due to wholesale loss of ECA and phenotypes due to accumulation of ECA intermediates [11].

**Fig. 1.**
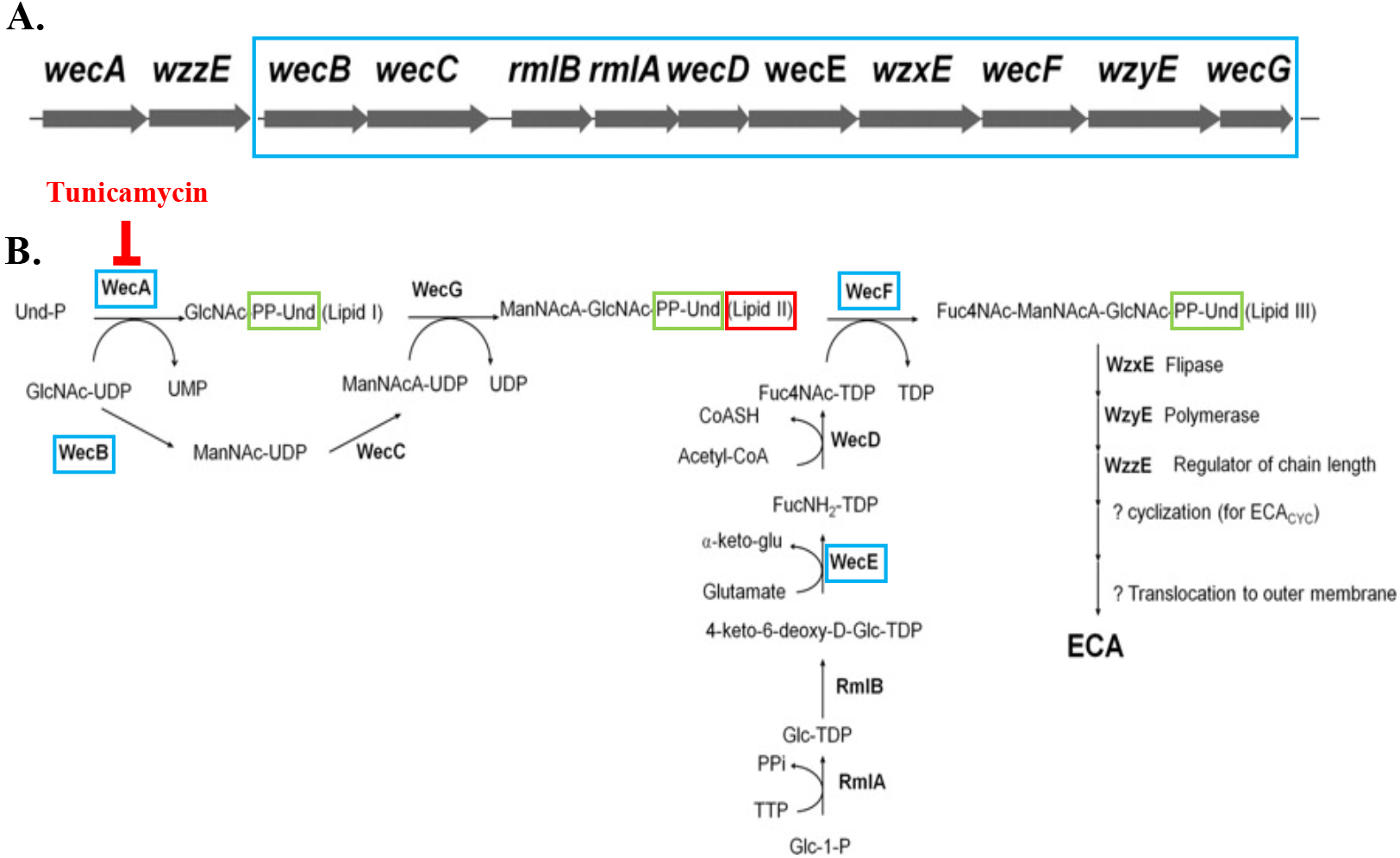
The ECA Biosynthesis Pathway. A) The *wec* operon. B) Enzymes involved in biosynthesis of ECA. Blue boxes indicate *wec* deletions made; *wecE/F* mutants lead to increased lipid II accumulation, while *wecA/B* deletions do not. Tunicamycin inhibits the WecA-mediated reaction so there is no buildup of lipid II intermediate (red) linked to the lipid carrier Und-PP (green). ECA is made up of repeating units of n-acetyl-D-glucosamine (GlcNAc), n-acetyl-D-mannosaminuronic acid (ManNAcA) and 4-acetamido-4,6-dideoxy-D-galactose (Fuc4NAc). Figure adapted from [11].

To verify the loss of ECA in these mutants, we used a dot-blot technique to determine the presence or absence of ECA. Ethanol-extracted ECA was dotted onto nitrocellulose and probed using *E. coli* O14 antisera, which has been previously shown to detect ECA [11]. All mutants in the ECA pathway showed severely decreased production of ECA as expected, while complementation in trans restored ECA production (Fig. 2).

**Fig. 2.**
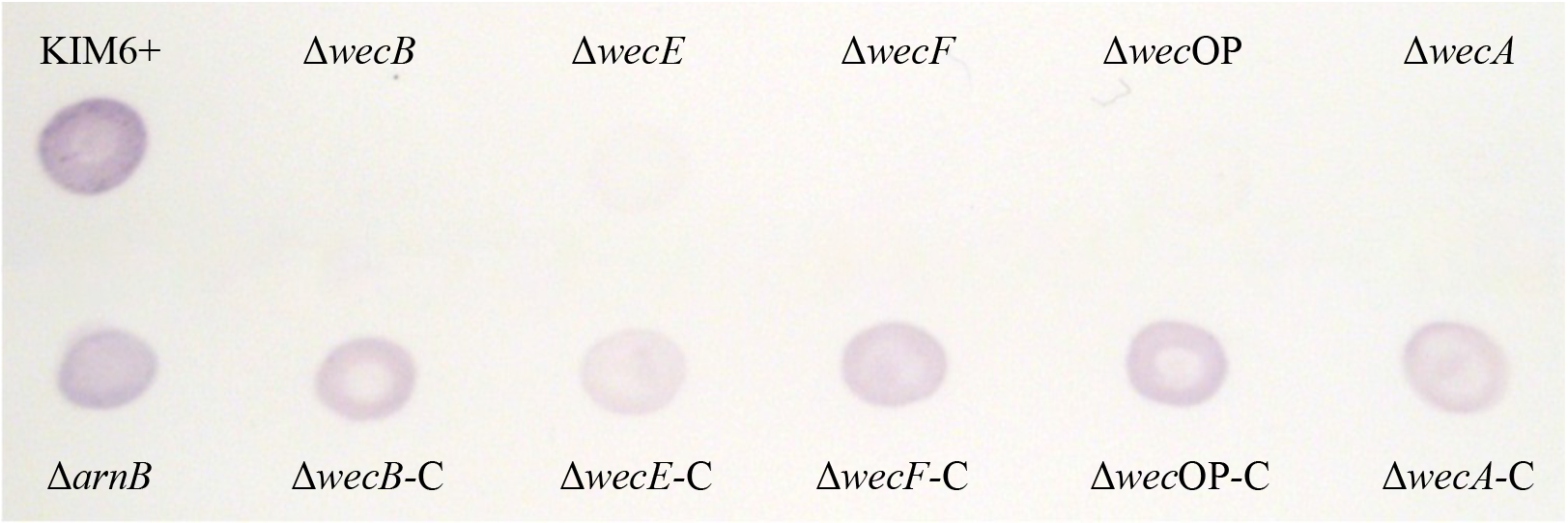
Dot blot for detection of ECA molecules. Purple color indicates the presence of ECA in cell extracts. (-C) indicates complementation with the appropriate gene(s).

### Sensitivity to PB is due to ECA-lipid II accumulation rather than lack of ECA

Deletion of genes encoding intermediate steps in the ECA biosynthesis pathway result in a buildup of the ECA-lipid II intermediate that cannot easily be recycled (Fig. 1), thereby interfering with normal cell envelope homeostasis by sequestering the limited amounts of the Und-P lipid carrier that is also required for other envelope maintenance pathways [7-9]. To examine how these ECA pathway deletions affect AMP resistance, we performed minimum inhibitory concentration (MIC) assays using PB with and without the addition of tunicamycin, which inhibits the first step in ECA biosynthesis (Fig. 1) [12]. Deletion of *wecA, wecB* or *wecOP*, which do not lead to buildup of ECA-lipid II intermediate, had MICs comparable to wild type KIM6+, while deletion of *wecE* or *wecF* both showed greatly increased sensitivity to PB (Table 1), suggesting sensitivity is due to issues from ECA-lipid II buildup rather than lack of ECA on the outer membrane. Furthermore, addition of tunicamycin to the culture media increased the resistance of the Δ*wecE/F* mutants to PB (Table 1), likely by increasing the availability of free Und-P to be used for other biosynthetic pathways, such as Ara4N.

**Table 1.** Polymyxin B Minimum Inhibitory Concentration.

| Strain | - Tunicamycin | + Tunicamycin |
| --- | --- | --- |
| KIM6+ | ≥256 | ≥256 |
| Δ*arnB* | 32 | 32 |
| Δ*wecB* | ≥256 | ≥256 |
| Δ*wecE* | 0.5 | ≥256 |
| Δ*wecF* | 0.5 | ≥256 |
| Δ*wecOP* | ≥256 | ≥256 |
| Δ*arnBwecE* | 0.125 | 0.125 |
| Δ*wecE* + *ispU* | 128 | ≥256 |
| Δ*wecF* + *ispU* | 128 | ≥256 |

To confirm that limitations in the pool of free Und-P is a key factor in the increased AMP sensitivity of these mutants, we overexpressed the gene responsible for production of Und-P in these strains to determine if resistance could be restored. Biosynthesis of the Und-P lipid carrier is dependent on *ispU* (*uppS*), which catalyzes the addition of eight isopentyl diphosphate molecules to farnesyl diphosphate, generating a 55-carbon chain Und-PP carrier which is then dephosphorylated to the Und-P active form [6]. Expression of an additional copy of *ispU* in the ECA mutants to increase the available pool of Und-P led to near wild-type levels of PB resistance in ECA-lipid II accumulating mutants (Table 1).

To determine if ECA pathway mutants showed alterations in their Ara4N-modified lipid A content, we performed MALDI-TOF mass spectrometry on lipid A extracted from ECA pathway mutants and control strains. We have previously used this method to examine changes in LPS modification in Ara4N biosynthetic pathway mutants [1]. The wild type KIM6+ strain displayed peaks corresponding to tetra (1HPO_3_)-, tetra (2HPO_3_)-, penta- and hexa-acylated lipid A species at *m/z* 1324,1404, 1586 and 1822, respectively, indicating unmodified lipid A, plus additional peaks corresponding to the Ara4N modification of each of these lipid A species (+131 *m/z*; Fig. 3). The lipid A spectra of ECA mutants that do not lead to ECA-lipid II buildup (Δ*wecB*, Δ*wecOP*) displayed similar Ara4N-modified lipid A profiles to wild type, with significant Ara4N modification of the major lipid A species at *m/z* 1455 and 1535. The Δ*wecE* and Δ*wecF* mutants showed decreased presence of these Ara4N peaks representing Ara4N modification (Fig. 3), which also correlates with their decreased levels of PB resistance.

**Fig. 3.**
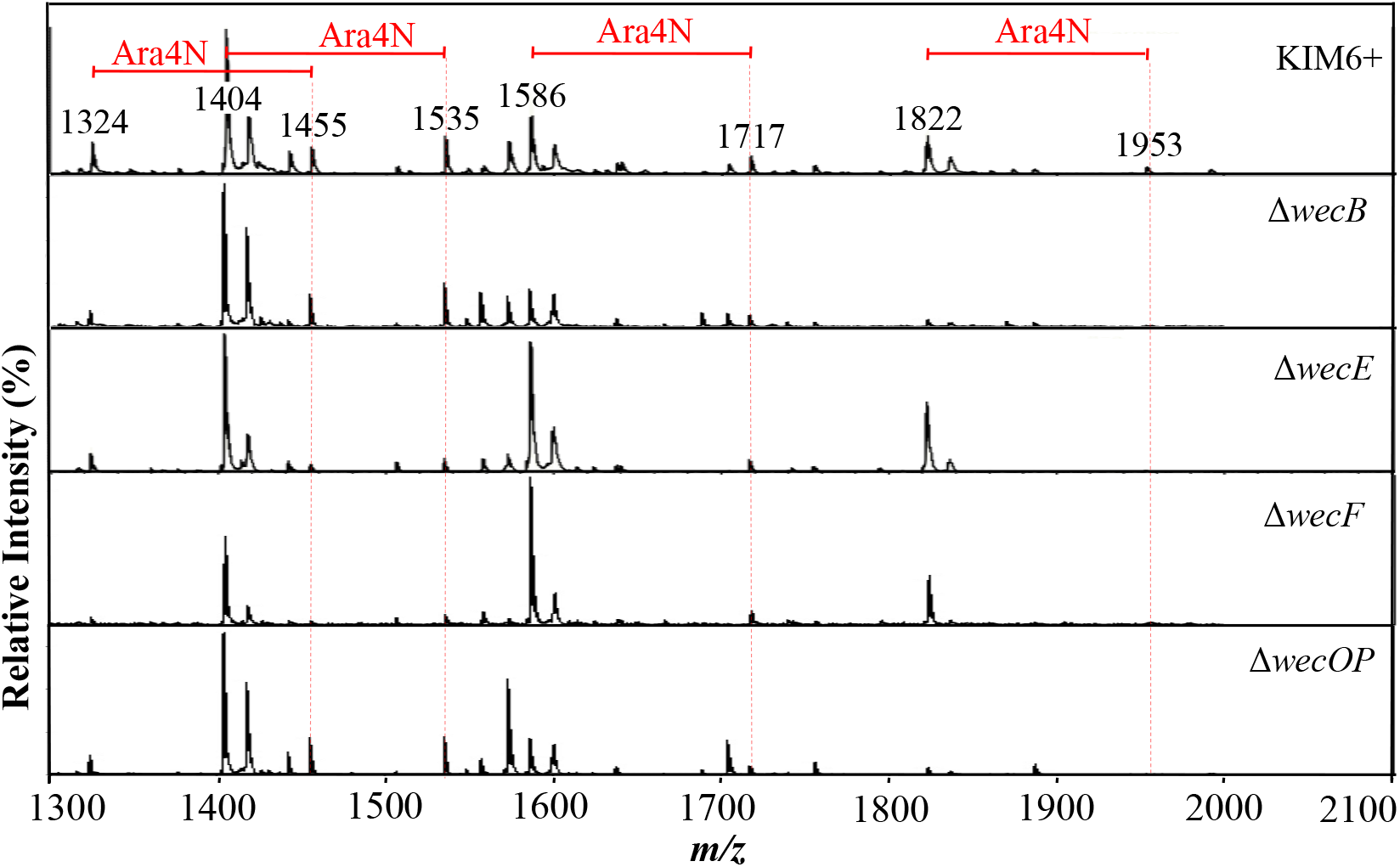
MALDI-TOF analysis of ECA pathway mutants. Mutants that accumulate ECA-lipid II intermediate show decreased Ara4N modification of major lipid A species. Ara4N modifications corresponding to lipid A species are indicated by red dotted lines (+131 *m/z)*.

### WecE mutants show altered cell morphology by microscopy

Biosynthetic pathways for outer membrane and cell wall components are known to compete for the pool of free Und-P lipid carrier in cells. In *E. coli*, sequestration of Und-P through “dead-end” pathways, such as ECA-lipid II or O-antigen intermediates, leads to morphological defects in cells, as well as membrane disruption and sensitivity to detergents and bile salts [9, 13]. To determine whether *Y. pestis* ECA pathway mutants share these characteristics we compared the morphology of mutant and wild type cells using microscopy. Cells were Gram-stained and images were processed using the ImageJ software to remove background and convert the cell images to black and white (Fig. 4). Individual cells were measured by counting the black pixels comprising each cell to give an estimate of cell size and the mean of 100 individual cells was calculated. Mutants lacking ECA (Δ*wecB*, Δ*wecOP*) had a mean cell size similar to KIM6+, while Δ*wecE* and Δ*wecF* mutants were skewed toward a larger mean cell size with obvious shape differences visible, being rounder and fatter than KIM6+ (Fig. 4). These differences suggest that sequestration of Und-P also affects other cell envelope component pathways in *Y. pestis*, including peptidoglycan biosynthesis, leading to cell growth and morphology changes.

**Fig. 4.**
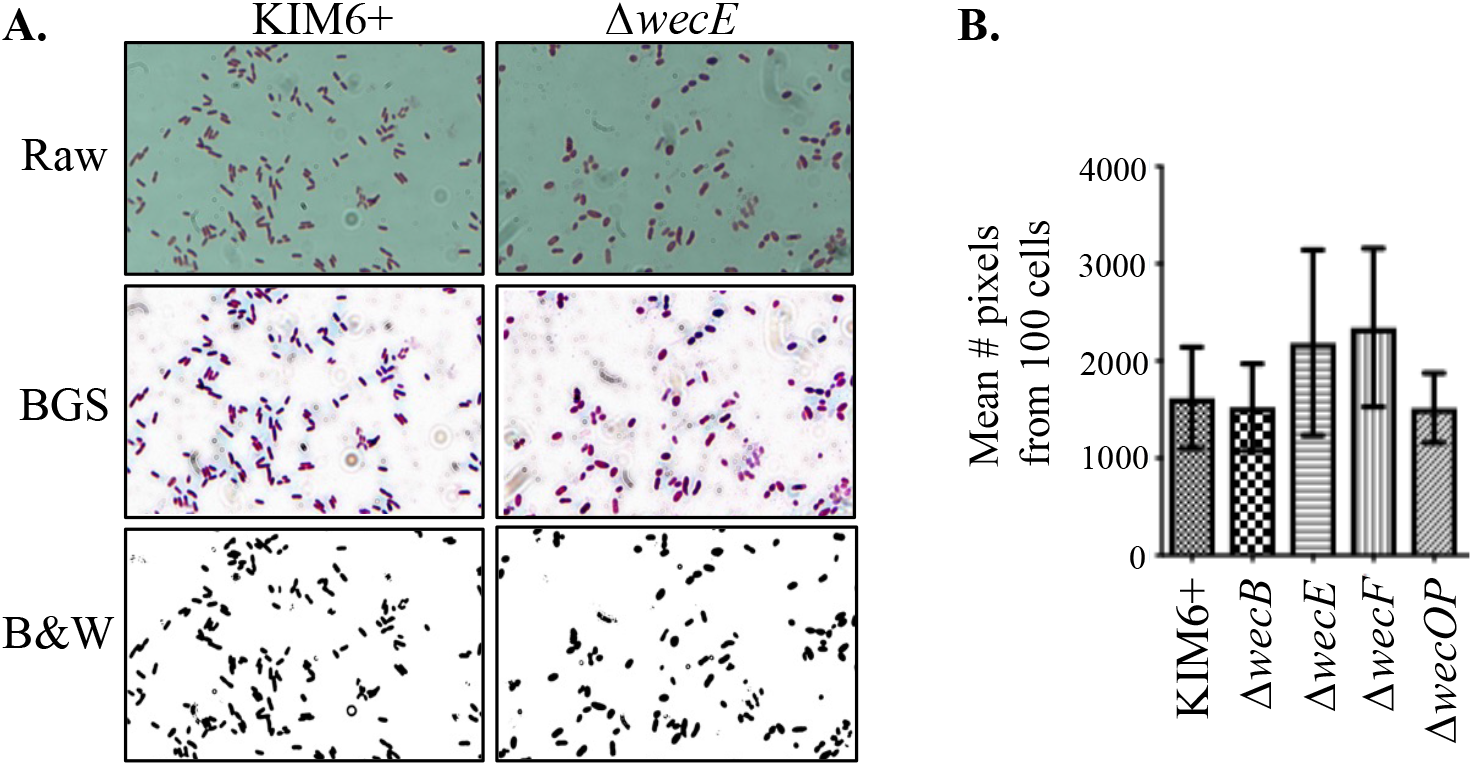
Mutants that accumulate ECA-lipid II show altered cell morphology. A) Microscopy of Gram-stained cells of KIM6+ and *ΔwecE*. Using ImageJ, raw image files were background subtracted (BGS) and converted to black and white (B&W). B) The mean number of pixels per cell was calculated for 100 individual cells from the B&W images of three independent experiments to estimate cell size.

### AMP-sensitive *wecE* mutants have decreased ability to colonize fleas

Defects in ECA biosynthesis have been shown to have numerous effects in different bacterial species, ranging from alterations in motility, to virulence, to maintaining outer membrane integrity [14-17]. To determine if lack of intact ECA, or decreased AMP resistance due to Und-P sequestration led to decreased flea colonization, we performed coinfection assays to examine the ability of Δ*wecE* and Δ*wecA* mutants to initiate successful infections in *X. cheopis* fleas in competition with wild type KIM6+ [18]. We coinfected fleas with bloodmeals containing each mutant strain, marked with a kanamycin (Km) cassette, and equivalent numbers of wild type KIM6+. Fed fleas were collected at 1 and 4 days post-infection to enable quantification of bacteria over the course of initial infection. Lysates from individual fleas were plated on media with and without Km and colonies were counted to determine the ratio of mutant to WT cells at each time point as compared to the input ratio of mutant to WT present in the infectious bloodmeal. The Δ*wecA*-Km mutant retained the ability to maintain a flea infection, while the Δ*wecE*-Km mutant was less able to mount a successful flea infection and showed a greatly decreased presence by day 4 (Fig. 5). This indicates that ECA itself does not play an important role in allowing *Y. pestis* to infect fleas, but mutants in the ECA pathway that might affect Und-P allocation can lead to decreased flea colonization ability, likely due to disruptions in the cell envelope and substantial loss of Ara4N, which increases AMP sensitivity.

**Fig. 5.**
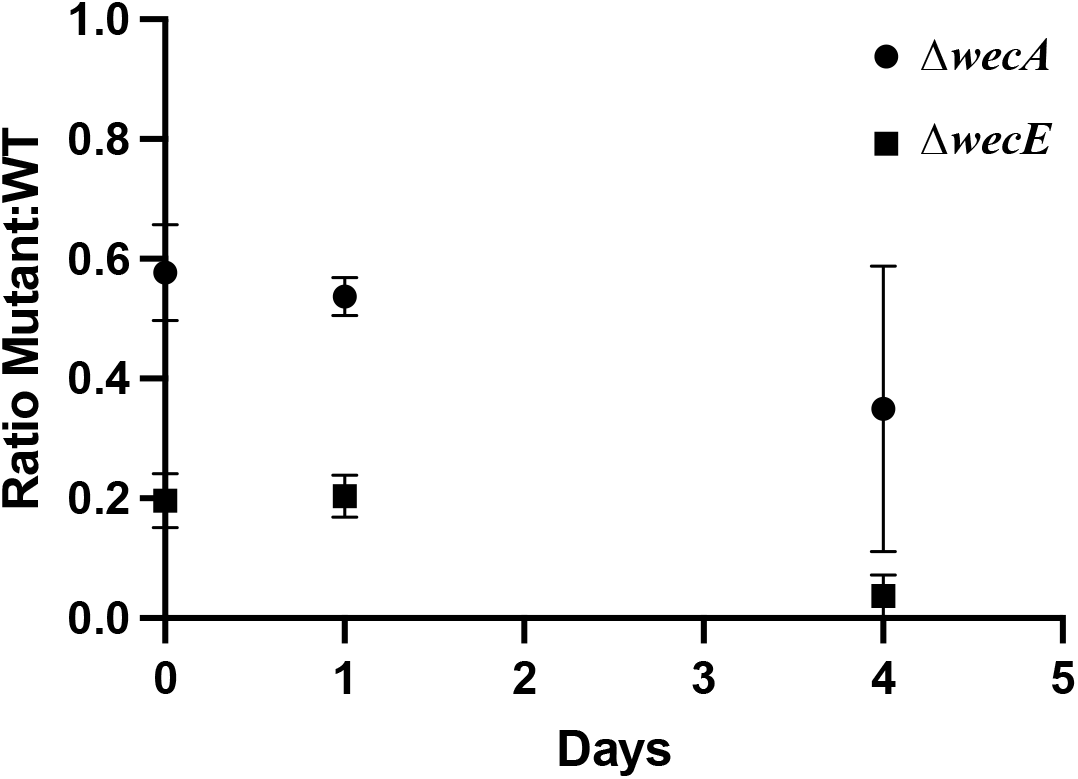
Ratio of mutant to wild type *Y. pestis* from individual fleas over time. Error bars show SD.

## DISCUSSION

In this study, mutations in the biosynthetic pathway for ECA were generated to determine how disruptions affect the AMP resistance of *Y. pestis. Yersinia pestis* KIM6+ is intrinsically resistant to a variety of AMPs, including Cecropin A and PB, and this resistance plays an important role in allowing it to mount a successful flea infection [1]. Previous research has demonstrated that *Y. pestis* mutants that cannot produce Ara4N-modified lipid A, a major determinant of AMP resistance, are highly sensitive to PB in vitro and cannot maintain wild type levels of infection in *X. cheopis* fleas. Deletion of the *wecE* gene, which is part of the ECA biosynthetic pathway, also resulted in PB sensitivity in *Y. pestis* [1]. To determine how biosynthesis of ECA affects AMP resistance, additional ECA deletion mutants were generated in *Y. pestis*, both in genes known to lead to an accumulation of ECA intermediates when disrupted, as well as genes that prevent initiation of ECA biosynthesis. Accumulation of ECA intermediates have been demonstrated to have far-reaching effects in bacterial cells, including changes in cell morphologies, sensitivities, and reactions to environmental stresses. Complete loss of ECA has also been implicated in the alteration of a variety of cellular processes, such as motility and virulence. However, assigning direct roles for ECA has proven difficult owing to the overlap of biosynthetic pathways of many cell envelope components that often require the same Und-P lipid carrier to enable biosynthesis. Limitations in Und-P availability can affect the biosynthesis of peptidoglycan, O-antigen, ECA, and Ara4N, all of which may alter the morphology and characteristics of a cell, making it difficult to determine the functions of specific genes and products of these pathways. Und-P limitations can occur when the carrier is allocated to a pathway from which it cannot be recycled, such as the Δ*wecE* mutant, in which the ECA-lipid II accumulates in the cell as ECA biosynthesis cannot be completed. However, by comparing ECA pathway mutants that accumulate intermediates (Δ*wecE/*Δ*wecF*) with those that do not (Δ*wecA/*Δ*wecB/*Δ*wecOP*), some insight can be gained into the roles of each.

Hypersensitivity to PB was identified only in ECA-lipid II-accumulating mutants, suggesting that intact ECA does not play a major role in AMP resistance in *Y. pestis*. Additionally, only the intermediate-accumulating ECA mutants showed obvious morphological defects by microscopy, suggesting abnormal cell wall structure. Sequestration of Und-P would be predicted to limit Ara4N biosynthesis and LPS modification, which was confirmed by MALDI-TOF for the Δ*wecE* and Δ*wecF* mutants, along with an expected increase in susceptibility to AMPs. The attenuation of the Δ*wecE* mutant in flea infection also supports this idea given that mutants lacking Ara4N are less able to successfully infect fleas.

Further investigation is needed to better understand how these altered, interconnected pathways affect cell growth and resistance and to identify their individual contributions. Although it is clear that the availability of Und-P plays an essential role in maintaining typical cell morphology and essential membrane barrier and resistance functions [7-9, 13], little is known about how cells allocate Und-P to the various pathways that require it. Further research into the production and distribution of this important glycan carrier is needed to determine how pathways obtain Und-P and how this process might vary between species. The results presented here are consistent with a model in which *Y. pestis* has a sufficiently limited pool of free Und-P such that sequestration of Und-P as ECA-lipid II prevents sufficient Ara4N biosynthesis, resulting in AMP hypersusceptibility. Given the central role it plays across multiple pathways affecting the cell envelope, Und-P is an appealing target for antimicrobial development. Further understanding of how it interacts with and affects these interconnected pathways could facilitate discovery of novel antibiotics, which in the face of rising antimicrobial resistance, are needed now more than ever.

## ACKNOWLEDGMENTS

We thank Kiara Held and Viveka Vadyvaloo (Washington State University) for assistance with flea coinfection assays and the University of Utah Health Sciences DNA and Peptide Core for DNA sequencing and primer synthesis.

This work was supported by a grant (R01AI130255) from the National Institutes of Allergy and Infectious Diseases.

