## Supplemental Table 1 and 2 for "Enterobacterial common antigen biosynthesis in *Yersinia pestis* is tied to antimicrobial peptide resistance"

**Table S1.** Bacterial strains and plasmids

| Strain or plasmid | Relevant characteristics/Usage | Reference or source |
|---|---|---|
| *Y. pestis* strains | | |
| KIM6+ | Parent strain; $pCD^-$, $pMT^+$, $pPCP^+$ | (1) |
| Δ*arnB* | In frame deletion of *arnB* in KIM6+ | (2) |
| Δ*wecE* | In frame deletion of *wecE* in KIM6+ | (2) |
| Δ*wecE*-C | Δ*wecE* complemented with pKLA-KM plus *wecE* gene | (2) |
| Δ*wecA* | In frame deletion of *wecA*in KIM6+ | This study |
| Δ*wecA*-C | Δ*wecA* complemented with pKLA-KM plus *wecA* gene | This study |
| Δ*wecB* | In frame deletion of *wecB* in KIM6+ | This study |
| Δ*wecB*-C | Δ*wecB* complemented with pKLA-KM plus *wecB* gene | This study |
| Δ*wecF* | In frame deletion of *wecF* in KIM6+ | This study |
| Δ*wecF*-C | Δ*wecF* complemented with pKLA-KM plus *wecF* gene | This study |
| Δ*wecOP* | In frame deletion of genes from *wecB* through *wecG* in KIM6+ | This study |
| Δ*wecOP-C* | Δ*wecOP* complemented with pKLA-KM plus *wecB-G* operon genes | This study |
| Δ*wecE*+*ispU* | **Δ***wecE* with PcysZK-*ispU* in pKLA-KM | This study |
| Δ*wecF*+*ispU* | **Δ***wecF* with PcysZK-*ispU* in pKLA-KM | This study |
| Δ*wecA*-Km | **Δ***wecA* with Km marker in Tn7 site for flea coinfection | This study |
| Δ*wecE*-Km | **Δ***wecE* with Km marker in Tn7 site for flea coinfection | This study |
| Plasmids | | |
| pKD46 | Lambda Red recombinase expression plasmid; $Ap^r$ | (3) |
| pKLA-KM | Complementation cloning plasmid; $Km^r$ | (2) |
| pFLP2 | Source of Flp recombinase; $Ap^r$ | (4) |
| pFKM1 | Km resistance cassette with flanking FRT sites; $Km^r$ | (5) |

**Table S2.** Primers

| Primer | Usage / Target | Primer Sequence (5’ → 3’) |
|---|---|---|
| wecA-Km-F | wecA λRed PCR | GGCATAGCAAGTGCTAATCTACATGCTAAGCTGTTGTATCT TTTTGCAGAGAGCGGTTAAACGTGAACCTACTCACTATGGC TCGAATTAGCTTCAAAAG |
| wecA-Km-R | wecA λRed PCR | AGTCGACATAGATTCTGGTTTCATCACTGCCTAAAAGCCTC TGGTTAAGATTCATGCTTATTTTTCGATGCCCGCCGCATAAT TGGGGATCTTGAAGTACC |
| wecB-Km-F | wecB/wecOP λRed PCR | GTATGAACTCGTATCGAGTCACGCACTGAGCGACAAGAAA TCGTTAATGGACGCATTGTGGCTCGAATTAGCTTCAAAAG |
| wecB-Km-R | wecB λRed PCR | TAACAGAAATAGTTTCAAAACTCATAGCGTCACCTGATGAT TCTTTAAAGCTTTAAGGATAATTGGGGATCTTGAAGTACC |
| wecF-UpR-Km | wecF λRed PCR | CTTCAGAGCGCTTTTGAAGCTAATTCGAGCCATTTTTTTCTA CTGTAAAGAGTGTTTACGG |
| wecF-DnF-Km | wecF λRed PCR | TTCGGAATAGGTACTTCAAGATCCCCAATTCAAGCGTTGGC TCTGGCTG |
| wecF-OutR | wecF λRed PCR/ Complementation | TTAAACTTAACCCGCCGAAA |
| wzxE-F | wecF λRed PCR | GGGTCAGCAGCATTTCAGAT |
| wecG-Km-R | wecOP λRed PCR | GCTGCGGCGTCAAGATAAGAAGGGGCGTGACTAAGAACAA AACACTACAGACGACCACTATAATAATAACCAACAAACTT AATTGGGGATCTTGAAGTACC |
| Km_int-F | Verifying λRed mutant | GCGAGTGATTTTGATGACGA |
| wecA-OutR | Verifying λRed mutant | TCAGTAATGGCTGTCGCACT |
| wecB-OutR3 | Verifying λRed mutant/ Complementation | CGCCTTCAACAGCAATCTTT |
| wecF-OutR3 | Verifying λRed mutant | GGCTGACGTAACCGTGTTTT |
| wecG-OutR | Verifying λRed mutant/ Complementation | GCCCGTATCGTTCCGTATTT |
| wecA-OutF | Complementation | GGCAGCCAACGTACATACAA |
| wecB-StartF | Complementation | GTAGCAATTGCGTTAATGGACGCATTGTGC |
| wecF-StartF | Complementation | GTCAGAATTCCGTAGACACTCAGCACCGTA |
